# The Microbial Metagenome and Tissue Composition in Mice with Microbiome-Induced Reductions in Bone Strength

**DOI:** 10.1101/562058

**Authors:** Jason D Guss, Erik Taylor, Zach Rouse, Sebastian Roubert, Catherine H Higgins, Corinne J Thomas, Shefford P Baker, Deepak Vashishth, Eve Donnelly, M Kyla Shea, Sarah L Booth, Rodrigo C Bicalho, Christopher J Hernandez

## Abstract

The genetic components of microbial species that inhabit the body are known collectively as the microbiome. Modifications to the microbiome have been implicated in disease processes throughout the body and have recently been shown to influence bone. Prior work has associated changes in the microbial taxonomy (phyla, class, species, etc.) in the gut with bone phenotypes but has provided limited information regarding mechanisms. With the goal of achieving a more mechanistic understanding of the effects of the microbiome on bone, we perform a metagenomic analysis of the gut microbiome that provides information on the functional capacity of the microbes (all microbial genes present) rather than only characterizing the microbial taxa. Male C57Bl/6 mice were subjected to disruption of the gut microbiota (ΔMicrobiome) using oral antibiotics (from 4-16 weeks of age) or remained untreated (n=6-7/group). Disruption of the gut microbiome in this manner has been shown to lead to reductions in tissue mechanical properties and whole bone strength in adulthood with only minor changes in bone geometry and density. ΔMicrobiome led to modifications in the abundance of microbial genes responsible for the synthesis of the bacterial cell wall and capsule; bacterially synthesized carbohydrates; and bacterially synthesized vitamins (B and K) (p <0.01). Follow up analysis focused on vitamin K, a factor that has previously been associated with bone health. The vitamin K content of the cecum, liver and kidneys was primarily microbe-derived forms of vitamin K (menaquinones) and was decreased by 32-66% in ΔMicrobiome mice compared to untreated animals (p < 0.01). Bone mineral crystallinity was decreased (p=0.01) was decreased in ΔMicrobiome mice (p < 0.001) and matrix carbonate-phosphoate ratio was increased. This study illustrates the use of metagenomic analysis to link the microbiome to bone phenotypes and implicates microbially synthesized vitamin-K as a regulator of bone matrix quality.

## INTRODUCTION

The gut microbiome consists of the genomic components, products, and microorganisms in the gastrointestinal tract ^(1)^. Changes in the constituents of the microbiome have been associated with a number of chronic diseases throughout the body including cardiovascular disease, obesity, diabetes, Alzheimer’s disease and arthritis ^(1)^. The effects of the microbiome on host physiology has resulted in considerable interest in the microbiome as a potential diagnostic or therapeutic target ^(2)^.

Recent studies have indicated that the microbiome can have a profound effect on bone: mice raised from birth in an environment completely absent of microbial life (germ-free) have altered long bone length and trabecular and cortical bone mass ^(3-5)^. Disruption of the gut microbiome using oral antibiotics lead to changes in trabecular and cortical bone mass and femoral geometry in mice ^(5-9)^. Manipulation of the gut microbiome with probiotics has been shown to reduce bone loss associated with estrogen depletion in mice ^(10-12)^ and recent studies have suggested a similar effect in humans ^(13)^. Together these findings implicate the microbiome as a contributor to bone mass and bone mineral density. However, bone mineral density does not completely explain fracture risk ^(14,15)^. The term “bone quality” is used to refer to characteristics of bone other than bone mineral density that influence bone strength and fracture risk ^(16)^. We recently demonstrated that disruption of the gut microbiome in mice led to reductions in femoral whole bone strength that could not be explained by changes in bone mass and geometry, indicating that modifications to the microbiome lead to impaired bone tissue quality (Fig 1A) ^(8)^.

**Figure 1.**
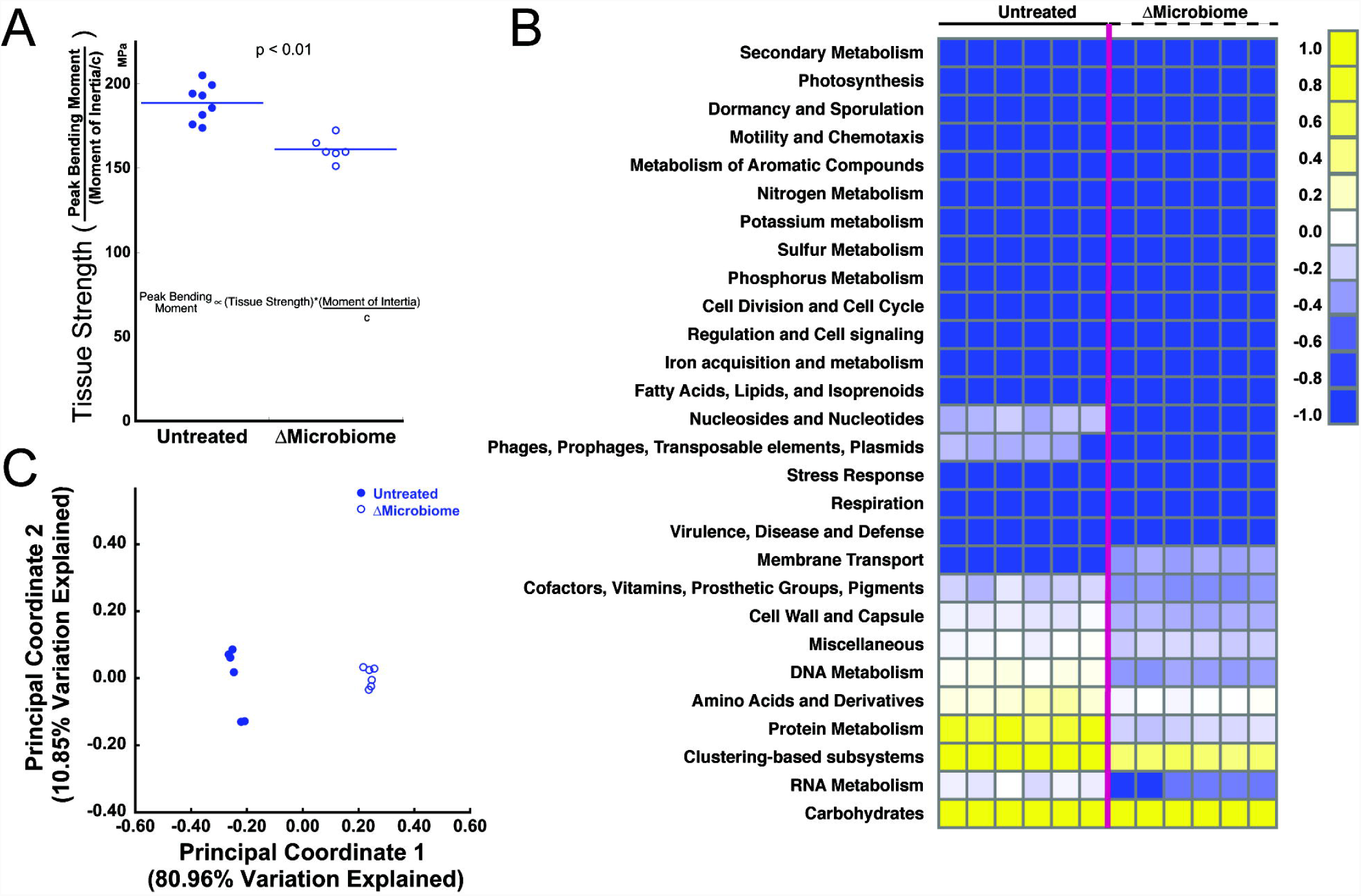
(A) Disruption of the gut microbiome (ΔMicrobiome) results in reductions in tissue strength assessed through three point bending of the mouse femur (figure adapted from ^(8)^). (B) A heatmap summarizing the metagenomic analysis of the fecal microbiota. Each column represents an individual animal (n=6 per group). (C) Principal coordinate analysis summarizes the differences in the functional capacity of the gut microbiota between the two groups.

To date, studies reporting an effect of the microbiome on bone have characterized the microbiome using sequencing of the bacterial 16S rRNA gene to determine the relative abundance of microbial taxa (phylum, class, order, etc.) ^(3,5,8,17)^. While phylogeny is useful for understanding the microbial community, more detailed sequencing often is required to identify molecular pathways that link the microbiome to host phenotype. Metagenomic sequencing involves analysis of the entire microbial genome and provides information on the functional capacity of the gut microbiome (i.e. which genes are present) ^(18,19)^. Metagenomic analysis is useful because many interactions between the microbiota and the host are a result of microbial functional capacity rather than microbial taxonomy.

Changes in the composition and structure of the organic or mineral composition of bone can lead to changes in both tissue- and whole bone mechanical performance ^(20-22)^. Bone tissue chemical composition can be assessed using Raman spectroscopy to determine: crystallinity (the size and stoichiometric perfection of the hydroxyapatite crystal lattice), mineral-to-matrix ratio (the extent of collagen mineralization and mineral content), and the carbonate-to-phosphate ratio (the extent of carbonate substitution into hydroxyapatite crystals). Additionally, nanoindentation can characterize compressive mechanical properties (hardness and reduced modulus) at the tissue-scale ^(20)^. To our knowledge, metagenomic analysis of the microbiome has not yet been used to understand the effects of the microbiome on bone. Additionally, although multiple studies report modifications in bone composition and nanomechanical properties in the context of bone quality and fracture risk ^(20,21,23-25)^, no previous studies have evaluated changes in bone tissue composition associated with changes in the gut microbiome.

The goal of this line of investigation is to determine how modifications to the gut microbiome can influence bone tissue quality. Using samples from a previously reported study including microbiome-induced changes in bone strength (Fig. 1A), we performed metagenomic analysis of the fecal microbiota as well as nanoscale chemical analysis of bone tissue. Specifically, we determined the changes in the fecal metagenome and bone tissue chemical composition and nanomechanical properties associated with microbiome-induced changes in bone tissue strength.

## 1.0 MATERIAL AND METHODS

### 2.1 Study design

Animal procedures were approved by Cornell University’s Institutional Animal Care and Use Committee. Mice from the C57BL/6J inbred strain were acquired (Jackson Laboratory, Bar Harbor, ME) and bred in conventional housing in our animal facility. Male mice were either treated to modify the gut microbiome (ΔMicrobiome) or untreated. Treated animals received broad-spectrum antibiotics (1.0 g/L ampicillin + 0.5 g/L neomycin) in their drinking water from weaning at 4 weeks of age until skeletal maturity (16 weeks of age) ^(26)^. Chronic antibiotics cause disruptions to the gut microbiome that are maintained over a prolonged time period ^(27)^. The oral bioavailability of these antibiotics is absent (neomycin) or low (ampicillin), thereby limiting extra-intestinal effects of dosing ^(26,28)^. Additionally, neomycin and ampicillin have never been associated with impaired bone growth, do not influence bone length, body mass or gut inflammation in these animals ^(8)^ and do not cause noticeable changes in serum calcium or vitamin D. Animals were housed in plastic cages filled with 1/4-inch corn cob bedding (The Andersons’ Lab Bedding, Maumee, Ohio), fed with standard laboratory chow (Teklad LM-485 Mouse/Rat Sterilizable Diet) and water *ad libitum*, and provided a cardboard refuge environmental enrichment hut (Ketchum Manufacturing, Brockville, Ontario). Animals were euthanized at 16 weeks of age. The right tibia, cecum, liver, kidney, and fecal samples were collected. Kidney and liver were stored at −20°C and cecum and fecal samples were stored at −80°C.

The study was performed using two cohorts of animals. One cohort of animals, described in a prior study ^(8)^, was used for metagenomic analysis and tissue chemical, nanomechanics and biochemistry (ΔMicrobiome n=7, untreated n=8). Tissue for some of the follow up biochemical analyses used samples from a second cohort of animals raised in our facility under identical conditions (untreated: n=6).

### 2.2 Metagenomic Analysis

Fecal samples collected one day prior to euthanasia were used for metagenomics analysis. Metagenomic analysis was performed on six samples per group (2 animals per cage). DNA was extracted (DNeasy PowerSoil DNA Isolation Kit, MO BIO Laboratories Inc., Carlsbad, CA) following manufacturer’s recommendations. The fecal pellet was added to the PowerBead tubes (Qiagen, Germantown, MD) and followed by a 10-minute vortex step. Following addition of Solution C1, to enhance cell lysis, samples were incubated at 70° C for 10 minutes and then subjected to a vortex step for 15 minutes using the MO BIO Vortex Adapter tube holder. Isolated DNA was quantified (Qubit dsDNA Broad Range Assay Kit, Life Technologies, Carlsbad, CA). Aliquots of DNA were normalized to the same concentration of 0.2 ng/ul of DNA per sample. A sequence library was prepared (Nextera XT DNA Library Preparation Kit, Illumina, San Diego, CA) to yield an average library size of 500 bp. Final equimolar libraries were sequenced (MiSeq reagent kit v3 on the MiSeq platform, Illumina, San Diego, CA) to generate 300 bp paired-end reads ^(29)^.

Metagenomic analyses were performed using MG-RAST (Metagenome Rapid Annotation using Subsystem Technology version 4.0.3) ^(30,31)^. In the MG-RAST analysis, the fragments of DNA in a sample are compared to protein, RNA, and subsystem databases. Functional annotation of sequences in the current study used the SEED subsystem ^(32)^. The functional abundance analysis was performed using a “Representative Hit Classification” approach with a maximum e-value of 1 × 10^−5^, minimum identity of 60%, and a minimum alignment length of 15 measured in amino acids for proteins and base pairs for RNA databases. The subsystems are grouped into hierarchical classifications ranging from the broadest functional category at “Level 1”, to more specific functional roles at “Level 2” and “Level 3”, and then to the most detailed category of “Function”. The data underwent a normalization and standardization process (within MG-RAST) to reduce inter-sample variability and to allow data to be more easily comparable. The normalized counts were calculated as: 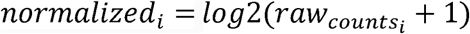. The standardized counts were calculated as: *standardized*_*i*_ = (*normalized*_*i*_ *-mean*(*normalized*_*i*_))/*stdev*(*normalized*_*i*_). Normalized counts are used as a measure of the abundance of genes that match a functional category. Differences in the abundance of genes in each of the functions were identified with α = 0.05 ^(33,34)^.

Principal coordinate analysis (PCoA) of the functional hierarchy based on the Bray-Curtis distance was performed to investigate overall functional diversity of the gut flora. Principal coordinate analysis reduces the dimensionality of a complex dataset with thousands of variables to a smaller number so the diversity between samples can be easily visualized in a two- or three-dimensional scatterplot ^(35)^. Each principal coordinate explains a percentage of the variation in the data set, with the first two principal components accounting for the most variation. PCoA was performed at subsystem Level 1, Level 2, and Level 3 hierarchies.

### 2.3 Raman Spectroscopy and Nanoindentation

The right tibiae were harvested and fixed in 10% neutral buffered formalin for 48 hours. Tibiae were then embedded undecalcified in methyl methacrylate and a single 2-mm-thick transverse section from the proximal metaphysis was collected using a diamond wafering saw (Buehler, Lake Bluff, Illinois). All sections were polished anhydrously on a Multiprep automatic polishing system (Allied High Tech, Rancho Dominguez, CA) at 30 RPM with a 200g sample load. Samples were polished with increasing grit silicon carbide polishing paper (800, 1200 grit) using ethylene glycol as a lubricant, and followed by a series of slurries of aluminum oxide powder (particle size of 3 μm, 1μm, and 0.1 μm) in ethylene glycol ^(36)^. The final root mean square (RMS) roughness of the surface was determined to be ∼35nm by measurement of ten 5 × 5-μm^2^ scans per sample with a surface profilometer (VKX Laser-Scanning Microscope; Keyence, Inc.).

A Raman imaging system (InVia Confocal Raman Microscope; Reinshaw Inc.) was used to collect spectra of the tibial cross sections by analyzing four different regions in each cross section (n=4/group). A total of 20 individual point spectra were collected across four quadrants of the cross section corresponding to 25%, and 75% of the cortical thickness with an additional three points collected 50 microns away from the midline of the cortex (forming a ‘+’ sign). The five spectra were averaged to determine a single representative measure per quadrant per sample. Spectra were collected over the range 720-1,820 cm^-1^ with a 785nm laser and a 50x long-working-distance objective (N.A.=0.55) collecting for 30s at 50% power with cosmic ray correction. Spectra first were normalized to the absorbance of PMMA at 813 cm^-1^ (MATLAB, MathWorks). Last, spectra were baseline-corrected to account for background fluorescence. The following Raman bands were evaluated: phosphate 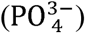 *v*_1_PO_4_ (integration area ∼930-980 cm^-1^) ^(37)^, amide III (integration area ∼1215-1300 cm^-1^) ^(37)^, and carbonate 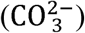 CO_3_ (integration area ∼1050-1100 cm^-1^) ^(37)^. From each spectrum the following measures were calculated: mineral-to-matrix ratio (determined as the area ratio of phosphate *v*_1_PO_4_ and amide III); carbonate substitution (measured as the area ratio of carbonate to phosphate *v*_1_PO_4_); and mineral crystallinity (measured as the inverse of the full-width-half-max of a Gaussian fit of the phosphate *v*_1_PO_4_ peak) ^(38)^.

Nanoindentation was performed on the same sections and regions analyzed by Raman spectroscopy. Nanoindentation arrays were performed using a Berkovich indenter tip (TI-900 Triboindenter, Bruker, Eden Prairie, MN) calibrated to a silica glass standard. Each array consisted of a 4 × 4 grid of indentations with a 30 second ramp load to *P*_*max*_ = 2500 μN, a 30 second hold to reach equilibrium, and a five-second elastic unloading. Indents were placed 15 μm away from each other to avoid mechanical interactions among indentations. Hardness (H) and reduced modulus (E_r_) were determined from the force vs. displacement curves of each indentation ^(39)^ using the following relations:

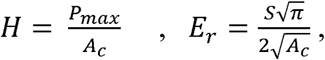

for which *S* is the contact stiffness (the slope of the load-displacement curve upon initial unloading) and A_c_ is the projected contact area of the indentation. The nominal contact depth of the indents in the bone samples was 260 nm.

### 2.4 Biochemical Analysis

Biochemical analyses of tissues were performed after receiving the results of the metagenomics analysis as a means of testing the functional significance of modifications to the microbial metagenome (n=6/group). Based on the metagenomics findings and bone biomechanical findings, the biochemical analysis focused on vitamin K. Vitamin K is a class of fat-soluble vitamers consisting of phylloquinone (PK, vitamin K_1_ in older literature) and the menaquinones (vitamin K_2_ in older literature). Menaquinones exist in ten known forms, identified by the length of the isoprenoid side chain of the molecule (labeled MK-*n* where n is the length of the side chain, MK4-MK13) ^(40)^. Phylloquinone and MK-4 are derived primarily from the diet. The remaining nine known forms of menaquinone are synthesized primarily by bacteria in the gut, although some bacterially-derived forms of vitamin K are found in fermented or cured food products ^(40)^. The cecum is an important site for microbial production of vitamin K ^(41)^. The liver and kidney are distant organs where vitamin K accumulates ^(42)^. Phylloquinone (PK) and menaquinone (MK-4-13) concentrations in the cecum, liver and kidney were measured by liquid chromatography/mass spectroscopy (LC/MS) ^(43)^. Detailed procedures for vitamin K extraction and sample purification are described elsewhere ^(43)^. The LC/MS system consists of an Agilent 6130 Quadrupole MSD with an atmospheric pressure chemical ionization (APCI) source connected to an Agilent series 1260 HPLC instrument (Agilent Technologies, Santa Clara, CA). Separations were completed using a reversed-phase C18 analytical column (Kinetex 2.6 μm, 150 mm × 3.0 mm; Phenomenex, Inc., Torrance, CA).

A major function of vitamin K in bone is carboxylation of Gla-containing proteins during bone formation. The most abundant Gla-containing protein in bone matrix is osteocalcin (also the most abundant non-collagenous protein in bone). To determine osteocalcin content, mouse humeri were dissected and wrapped in PBS soaked gauze. The tested mouse humeri were homogenized in 600 μl of extraction buffer containing 0.05M EDTA, 4M guanidine chloride and 30mM Tris-HCl (Omni BeadRuptor 24, Omni International, Atlanta, GA). After homogenization, the solution was centrifuged at 13000 rpm for 15 minutes to eliminate remaining mineral debris from the supernatant. The supernatant was dialyzed against 1x PBS and 5mM EDTA for two days to eliminate denaturant. Extracted bone protein concentrations of the dialyzed solutions were assessed using a Pierce™ Coomassie Plus (Bradford) Assay Kit. The extracts then were serially diluted 1000-fold in PBS for use with the LSBio Mouse OC ELISA kit, which has a working range of 0.156-10 ng/mL. The OC quantification ELISA was performed as per manufacturer protocol. Osteocalcin content was assessed in 4-5 animals per group.

### 2.5 Statistical Treatment

Group differences between nanoindentation measures, metagenome sequence abundances, vitamin K levels, and osteocalcin content were determined using a one-way ANOVA α=0.05 (JMP Pro 9.0.0). Differences in Raman measures between groups were determined using a generalized least squares model (GLM) to account for the effect of quadrant.

## 3.0 RESULTS

### 3.1 Metagenome Functional Analysis

The functional capacities of the gut microbiome differed among groups (Fig 1B). Disruption of the gut microbiome caused drastic changes in the functional capacity of the gut microbiome, as indicated by distinct clusters in the principal coordinate analysis (Fig 1C). Metagenomics findings indicated differences in the abundance of genes associated with vitamin biosynthesis, carbohydrate function and cell and cell capsule synthesis.

Pathways related to the synthesis of vitamin B and vitamin K were altered by disruption of the gut microbiome. Mice with a disrupted gut microbiome had lower normalized counts for genes associated with the synthesis of vitamin B2, B6, and B7 compared to untreated mice (Fig 2A), but had greater normalized counts for genes involved in the synthesis of vitamin B9 and K. Further investigation identified differential presence of multiple genes involved in menaquinone biosynthesis (Fig 2B): normalized counts for MenH, MenF, and MenE genes are greater in ΔMicrobiome mice and the abundance of MenB, MenC and MenG genes was less in ΔMicrobiome mice than in untreated mice.

**Figure 2.**
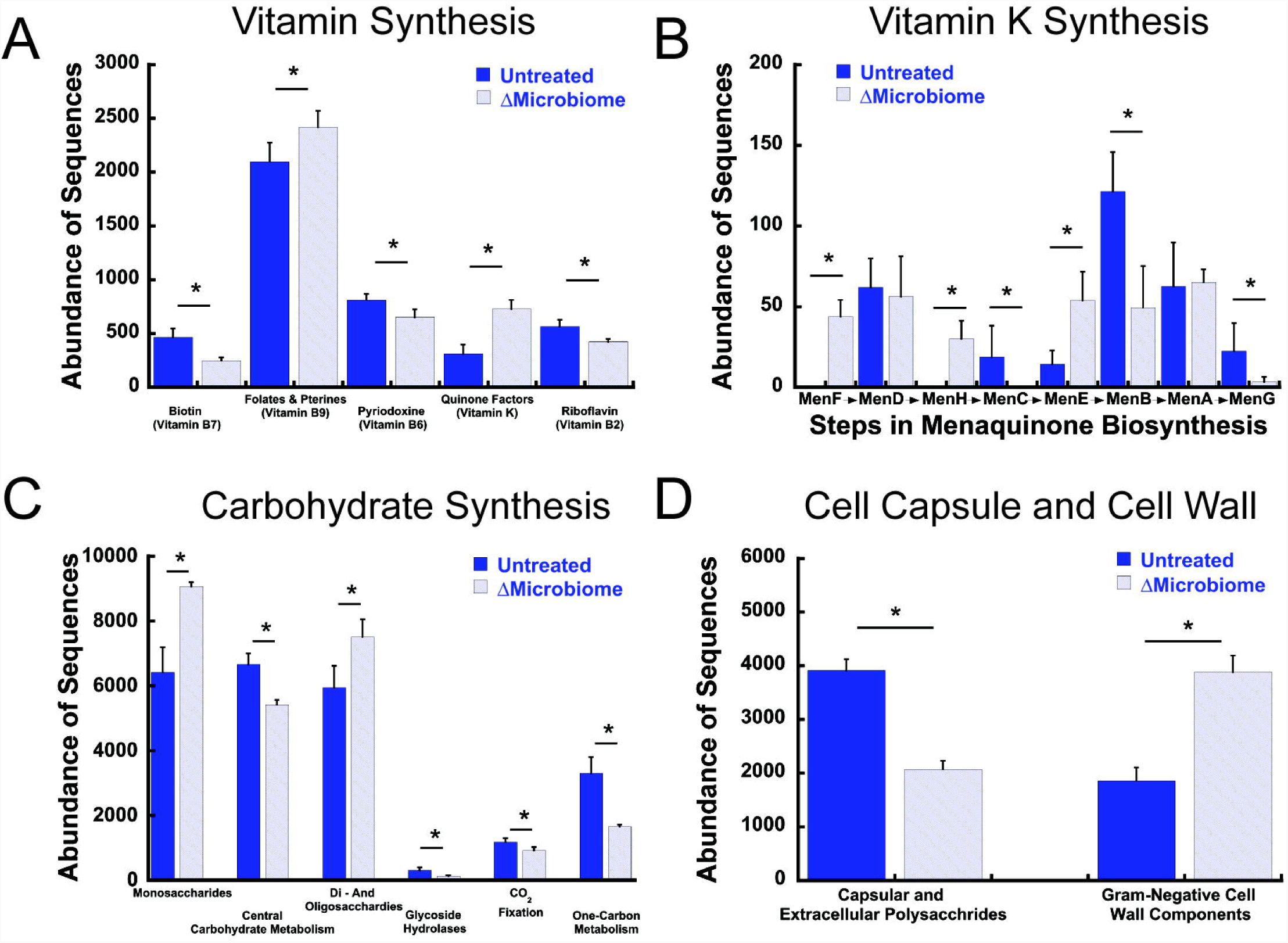
The relative abundance of genes associated with key pathways for (A) vitamin synthesis, (B) vitamin K synthesis (shown in the order of synthesis), (C) carbohydrates, and (D) bacterial cell wall and capsule components are altered in ΔMicrobiome mice (n=6/group). * - p < 0.002.

The overall functional capacity and the abundance of genes for six of eight carbohydrate functional categories were altered by ΔMicrobiome (Fig 2C). No differences in the overall abundance of fermentation genes were detected. The abundance of genes related to the cell wall and cell capsule differed among groups (Fig 2D). Normalized counts for genes for capsular and extracellular polysaccharides were less abundant in mice with a disrupted gut microbiome than in untreated mice (p < 0.01). Disruption of the gut microbiome led to increased abundance of genes associated with Gram-negative cell wall components.

### 3.2 Raman Spectroscopy and Nanoindentation

Bone tissue crystallinity, carbonate substitution and mineral to matrix ratio varied among quadrants (p < 0.05, Fig. 3). After accounting for variation among quandrants, disruption of the gut microbiome was associated with decreased crystallinity (p=0.01, average difference 0.00056), increased carbonate substitution (p < 0.001, average difference 0.0113) and no detectable differences in mineral:matrix (Fig. 3B-D).

**Figure 3.**
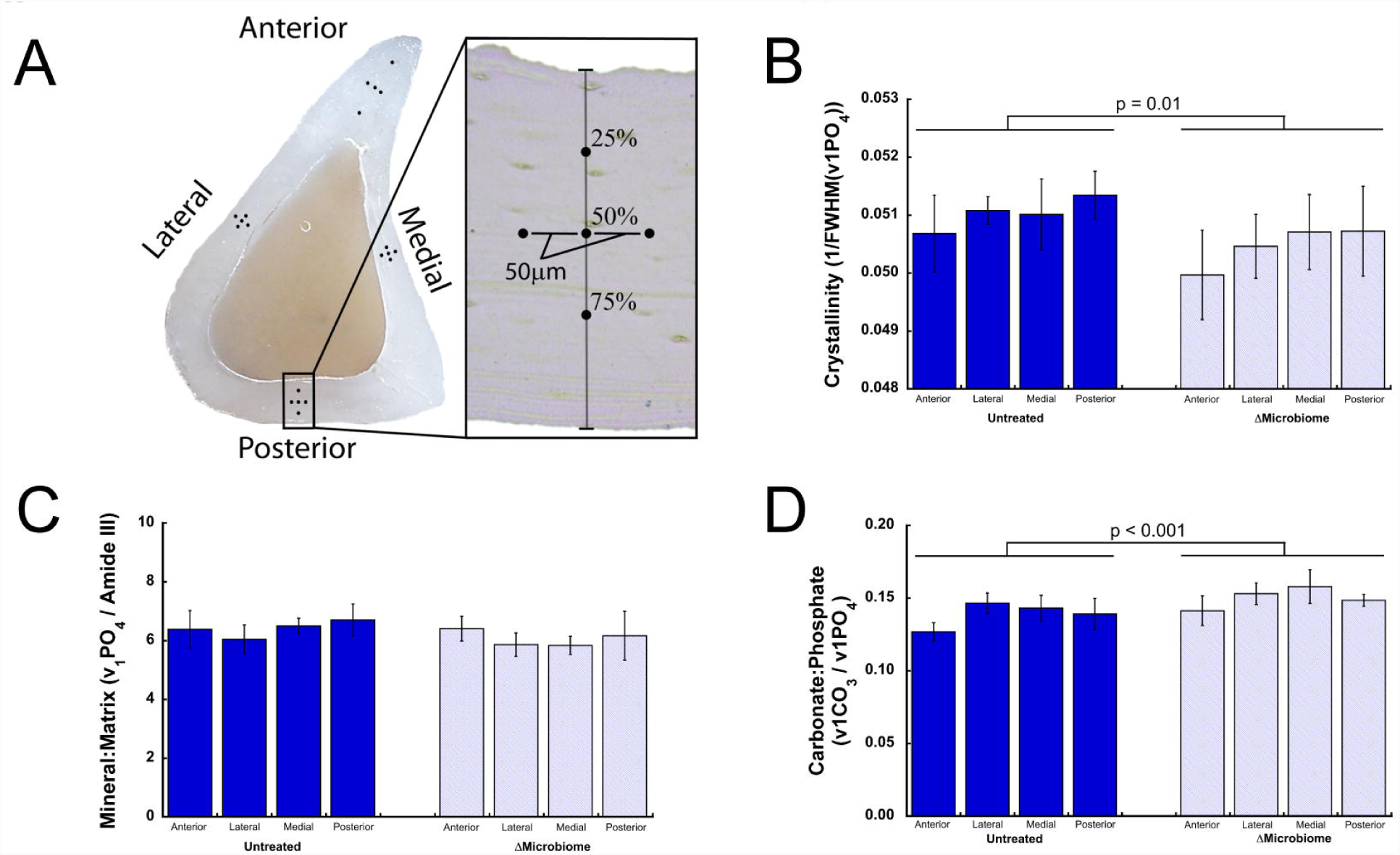
(A) Five Raman point spectra were collected in each of the four anatomical quadrants of a tibial diaphysis cross section. The average of the five spectra within each quadrant was determined. ΔMicrobiome was associated with reduced (B) crystallinity, no noticeable differences in (C) mineral:matrix ratio and an increase in (D) carbonate substitution after accounting for variation among quadrants (n=4 specimens/group, error bars indicate SD).

Reduced modulus measured using nanoindentation was similar among groups (Supplemental Fig 4A; untreated: 30.8 GPa ± 1.06; ΔMicrobiome: 30.4 GPa ± 1.20, mean ± SD). Hardness was similar among groups (Supplemental Fig. 1; untreated: 1.08 GPa ± 0.07; ΔMicrobiome: 1.09 GPa ± 0.04).

**Figure 4.**
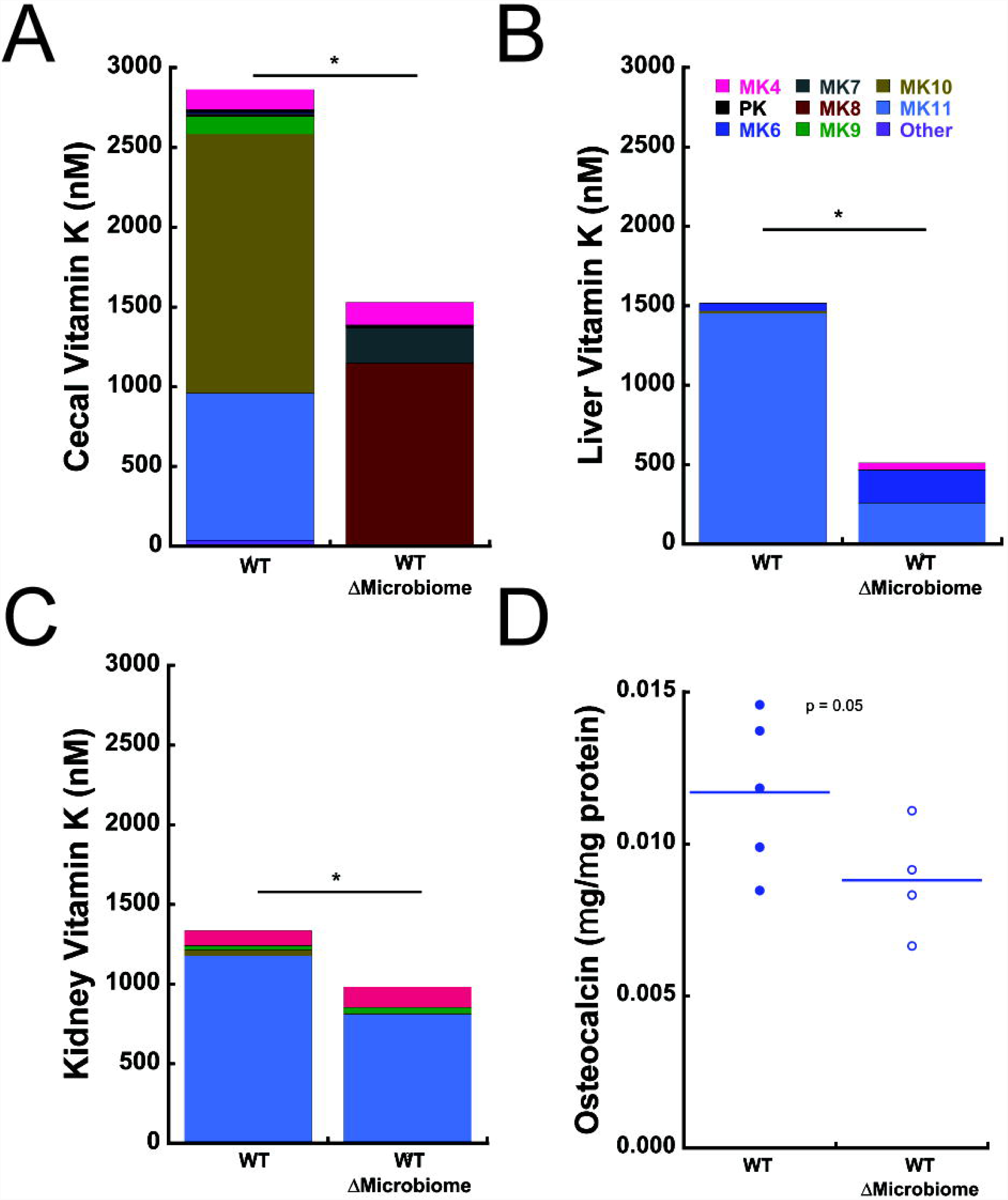
Vitamin K content was altered by disruption of the gut microbiome in the (A) Cecum, (B) Liver and (C) Kidney. n=6/group * indicates p < 0.001.(D) Matrix bound osteocalcin trended toward reduction in ΔMicrobiome mice (p=0.05).

### 3.5 Biochemical Analysis

Vitamin K content in the cecum, liver, and kidney primarily consisted of microbe-derived menaquinones; on average, the microbe-derived menaquinones (MK5-13) accounted for 83.3% to 99.9% of the total vitamin K content (Fig 4A-C). Total cecal vitamin K content was lower in ΔMicrobiome mice compared to untreated mice (Fig 4A). Total liver vitamin K content was lower in ΔMicrobiome compared to untreated mice (Fig 4B). Kidney vitamin K content was also decreased in ΔMicrobiome mice (Fig 4C). Mean matrix-bound osteocalcin concentration was reduced in ΔMicrobiome mice (p = 0.05, Fig. 4D).

## 4.0 Discussion

This study provides the first report of the metagenomic components of the microbiome in a situation of altered bone. The metagenomic analysis identified differences among groups in terms of the abundance of genes related to vitamin synthesis, cell wall and capsule synthesis, and carbohydrate synthesis. The observed differences in the abundance of genes associated with vitamin synthesis led to follow up biochemical analyses focused on vitamin K, a factor that has long been associated with bone health ^(44,45)^. Biochemical analysis confirmed reduced concentrations of vitamin K in the cecum, liver and kidney associated with disruption of the gut microbiota – an effect dominated by reductions in the concentrations of forms of vitamin K generated by microbes (menaquinones 5-13), supporting a potential link between vitamin K produced by the gut microbiota and bone tissue quality.

The current study provides a metagenomic analysis as a means of identify potential mechanistic relationships between disruption of the gut microbiome and impaired bone tissue strength (Fig. 1A). The gut microbiome may influence bone tissue through three general mechanisms: 1) regulation of nutrient absorption and microbe-derived vitamins; 2) regulation of the immune system; and 3) translocation of inflammatory bacterial products across the gut barrier ^(46)^. While regulation of the immune system and translocation of inflammatory bacterial products can lead to changes in bone resorption, bone formation and bone mass ^(47)^, these mechanisms are only known to regulate bone matrix quality only by modifying tissue age, a factor that does not vary much in mice at 16 weeks of age. In contrast, vitamins produced by the gut microbiota can influence bone tissue. In particular, vitamin K is produced by the gut microbiota and has long been associated with bone health ^(44,45)^.

Together with prior work, our findings provide preliminary support for a potential link between the microbiome and bone tissue quality that is mediated by microbiome-derived vitamin K. Although vitamin K may influence bone tissue quality in multiple ways, the best understood mechanism is γ carboxylation of gamma-carboxyglutamic (Gla-) containing proteins ^(48)^. Vitamin K-dependent γ carboxylation is required for proper binding of Gla-containing proteins to bone tissue ^(48,49)^. Bone contains many vitamin K-dependent proteins, however, the vitamin K-dependent protein osteocalcin is the most abundant non-collagenous protein in bone tissue and is known to influence bone tissue mechanical properties ^(50,51)^. Interestingly, our biochemical analysis suggests that ΔMicrobiome may lead to reductions in matrix-bound osteocalcin (p = 0.05). When present in bone tissue, non-collagenous proteins such as osteocalcin can regulate and direct the formation and size of collagen fibrils, as well as mineralization and crystal nucleation, leading to changes in crystallinity ^(52-56)^. Crystallinity is descriptive of the size, perfection, and maturity of hydroxyapatite crystals and reductions in matrix crystallinity are associated with reduced bone tissue strength ^(57)^. Osteocalcin-deficient mice have decreased crystallinity ^(58)^ and decreased bone tissue strength ^(59)^. Similarly, we found ΔMicrobiome to lead to reduced crystallinity in this cohort of animals with impaired tissue strength (Fig. 1A). Together these findings implicate vitamin K as a potential link between the microbiome and bone tissue strength, but does not prove causation. Hence, we cannot ignore the potential contribution of other mechanisms through which the microbiome may mediate bone.

In addition to identifying differences in vitamin synthesis, the metagenomic analysis also observed significant changes in the abundance of genes associated with cell wall and capsule synthesis and carbohydrate synthesis. We attribute the differences in abundance of cell wall and capsule genes with changes in the taxonomic components of the gut flora in this cohort. Specifically, our prior taxonomic analysis associated ΔMicrobiome with increases in the abundance of organisms from the Gram negative phyla *Proteobacteria* in this cohort ^(8)^, which is consistent with the increase in the abundance of genes associated with production of Gram negative cell capsule components. In contrast, the observed changes in abundance of genes associated with carbohydrate synthesis is not as easily explained by taxonomy. These genes can influence the production of molecules such as short chain fatty acids that have been associated with changes in bone formation and remodeling ^(5)^, although a mechanism through which these proteins might influence bone tissue quality has not yet been proposed.

The changes in bone tissue chemistry observed here are consistent with modifications in whole bone mechanical performance reported previously for this cohort (Fig. 1A). Reduced crystallinity has previously been correlated with reduced bone tissue strength and/or stiffness in humans and animals ^(57,60,61)^. Although this cohort shows reduced tissue strength assessed in bending, we did not observe differences in nanoindentation-derived elastic modulus or hardness, a finding we attribute to the fact that nanoindentation describes primarily compressive properties of bone tissue while bending strength is determined primarily by failure properties in tension ^(23)^.

Several strengths in the study are worth noting. To our knowledge the current study is the first to associate changes in the gut flora metagenomic constituents with bone. Previous studies have reported changes in phylogenetic profile using 16S rRNA sequencing ^(3,5,8,17)^. Because many different microbes have the same functional capacity, a shift in the microbial taxa may not represent differences in the functions of the microbiota. By providing the functional capacity, metagenomic analysis provides more information about potential links between the microbiome and bone. Second, to our knowledge, the current study is the first to evaluate how alterations to the gut microbiome can influence bone tissue composition and material properties. Most previous studies have focused on how the gut microbiome can influence bone microstructure and bone remodeling, but have not reported bone chemistry and nanoscale properties. Lastly, the vitamin K assays allowed for the differentiation between dietary and microbe-derived forms of vitamin K. Previous studies evaluating vitamin K and bone phenotype in rodents have been restricted to phylloquinone or only one menquinone ^(62-64)^.

Despite the strengths of the current study, a few limitations must be considered when interpreting the findings. First, with regard to the metagenomic analysis, the current study was hypothesis-generating and, as molecules of interest were not known a priori, it was not possible to design the study with statistical power for all follow up biochemical assays (matrix osteocalcin in particular). Despite this limitation, the reductions in cecal and kidney vitamin K and bone tissue crystallinity in ΔMicrobiome mice and the trend toward reduced osteocalcin content were all consistent with a potential microbiome – vitamin K - matrix osteocalcin mechanism. However, the effects of vitamin K may be a result of other vitamin K-dependent molecules in bone tissue (matrix Gla protein, etc.) or other ligands of vitamin K in the body (the pregnane X receptor, for example ^(65)^). Additionally, although Raman spectroscopy is useful for examining chemical composition, other modifications in tissue composition may be present that are not well described by Raman spectroscopy. Third, the biochemical analysis focused only on vitamin K in the cecum, liver, and kidney. Future studies will require a more comprehensive testing of other key potential factors such as vitamin B, circulating MAMPs such as LPS, and intestinal short-chain fatty acids.

In conclusion, we find that disruptions to the gut microbiome that lead to impaired bone tissue mechanical properties also lead to drastic shifts in the overall functional capacity of the gut microbiome. We observed shifts in functional capacity of the gut microbiota that were associated with changes in bone mineral crystallinity, the degree of carbonate substitution, and concentrations of microbially-derived forms of vitamin K in the body. Together our findings support the use of metagenomics for a microbiome analysis, and provide preliminary evidence for a mechanism in which production of vitamin K by the gut flora may influence downstream pathways responsible for bone tissue composition and structure.

## 5.0 Acknowledgments

This publication was supported in part by the National Institute of Arthritis and Musculoskeletal and Skin Diseases of the National Institutes of Health (U.S) under Award Number AR068061 and by the Department of Defense Congressionally Directed Medical Research Programs under Award Number W81XWH-15-1-0239. The content of the work is solely the responsibility of the authors and does not necessarily represent the official views of the National Institutes of Health or the Department of Defense. Additional funding was obtained from the USDA ARS Cooperative Agreement 58 −1950 −7 −707. Any opinions, findings, or conclusion expressed in this publication are those of the authors and do not necessarily reflect the view of the US Department of Agriculture.

Authors’ roles: Conceived and designed the experiments: JDG, CJH, RCB, SLB, MKS, DV, SPB, ED. Performed the experiments: JDG, SR, ZR, CHH, CJT, MKS. Analyzed data: JDG, CJH, ET. Wrote and Revised Manuscript: JDG, CJH. Critical revision and final approval of the manuscript: All authors.

We would also like to acknowledge Marjolein CH van der Meulen for her feedback in the preparation of the manuscript.

## References

1. Knight R, Callewaert C, Marotz C, Hyde ER, Debelius JW, McDonald D, Sogin ML 2017 The Microbiome and Human Biology. Annu Rev Genomics Hum Genet.

2. Gilbert JA, Blaser MJ, Caporaso JG, Jansson JK, Lynch SV, Knight R 2018 Current understanding of the human microbiome. Nat Med 24(4):392–400.

3. Schwarzer M, Makki K, Storelli G, Machuca-Gayet I, Srutkova D, Hermanova P, Martino ME, Balmand S, Hudcovic T, Heddi A, Rieusset J, Kozakova H, Vidal H, Leulier F 2016 Lactobacillus plantarum strain maintains growth of infant mice during chronic undernutrition. Science 351(6275):854–7.

4. Sjogren K, Engdahl C, Henning P, Lerner UH, Tremaroli V, Lagerquist MK, Backhed F, Ohlsson C 2012 The gut microbiota regulates bone mass in mice. J Bone Miner Res 27(6):1357–67.

5. Yan J, Herzog JW, Tsang K, Brennan CA, Bower MA, Garrett WS, Sartor BR, Aliprantis AO, Charles JF 2016 Gut microbiota induce IGF-1 and promote bone formation and growth. Proc Natl Acad Sci U S A.

6. Cho I, Yamanishi S, Cox L, Methe BA, Zavadil J, Li K, Gao Z, Mahana D, Raju K, Teitler I, Li H, Alekseyenko AV, Blaser MJ 2012 Antibiotics in early life alter the murine colonic microbiome and adiposity. Nature 488(7413):621–6.

7. Cox LM, Yamanishi S, Sohn J, Alekseyenko AV, Leung JM, Cho I, Kim SG, Li H, Gao Z, Mahana D, Zarate Rodriguez JG, Rogers AB, Robine N, Loke P, Blaser MJ 2014 Altering the intestinal microbiota during a critical developmental window has lasting metabolic consequences. Cell 158(4):705–21.

8. Guss JD, Horsfield MW, Fontenele FF, Sandoval TN, Luna M, Apoorva F, Lima SF, Bicalho RC, Singh A, Ley RE, van der Meulen MC, Goldring SR, Hernandez CJ 2017 Alterations to the Gut Microbiome Impair Bone Strength and Tissue Material Properties. J Bone Miner Res 32(6):1343–1353.

9. Nobel YR, Cox LM, Kirigin FF, Bokulich NA, Yamanishi S, Teitler I, Chung J, Sohn J, Barber CM, Goldfarb DS, Raju K, Abubucker S, Zhou Y, Ruiz VE, Li H, Mitreva M, Alekseyenko AV, Weinstock GM, Sodergren E, Blaser MJ 2015 Metabolic and metagenomic outcomes from early-life pulsed antibiotic treatment. Nat Commun 6:7486.

10. Britton RA, Irwin R, Quach D, Schaefer L, Zhang J, Lee T, Parameswaran N, McCabe LR 2014 Probiotic L. reuteri treatment prevents bone loss in a menopausal ovariectomized mouse model. J Cell Physiol 229(11):1822–30.

11. Li JY, Chassaing B, Tyagi AM, Vaccaro C, Luo T, Adams J, Darby TM, Weitzmann MN, Mulle JG, Gewirtz AT, Jones RM, Pacifici R 2016 Sex steroid deficiency-associated bone loss is microbiota dependent and prevented by probiotics. J Clin Invest.

12. McCabe LR, Parameswaran N 2018 Advances in Probiotic Regulation of Bone and Mineral Metabolism. Calcif Tissue Int 102(4):480–488.

13. Ohlsson C, Curiac D, Sjogren K, Jansson P 2018 Probiotic treatment using a mix of three lactobacillus strains protects against lumbar spine bone loss in health early postmenopausal women American Society for Bone and Mineral Research, Montreal, Canada, pp 1071.

14. Hillier TA, Stone KL, Bauer DC, et al. 2007 Evaluating the value of repeat bone mineral density measurement and prediction of fractures in older women: The study of osteoporotic fractures. Arch Intern Med 167(2):155–160.

15. Schuit SCE, van der Klift M, Weel AEAM, de Laet CEDH, Burger H, Seeman E, Hofman A, Uitterlinden AG, van Leeuwen JPTM, Pols HAP Fracture incidence and association with bone mineral density in elderly men and women: the Rotterdam Study. Bone 34(1):195–202.

16. Hernandez CJ, Keaveny TM 2006 A biomechanical perspective on bone quality. Bone 39(6):1173–81.

17. Blanton LV, Charbonneau MR, Salih T, Barratt MJ, Venkatesh S, Ilkaveya O, Subramanian S, Manary MJ, Trehan I, Jorgensen JM, Fan YM, Henrissat B, Leyn SA, Rodionov DA, Osterman AL, Maleta KM, Newgard CB, Ashorn P, Dewey KG, Gordon JI 2016 Gut bacteria that prevent growth impairments transmitted by microbiota from malnourished children. Science 351(6275).

18. Sharpton TJ 2014 An introduction to the analysis of shotgun metagenomic data. Front Plant Sci 5:209.

19. Thomas T, Gilbert J, Meyer F 2012 Metagenomics - a guide from sampling to data analysis. Microb Inform Exp 2:3–3.

20. Gourion-Arsiquaud S, Faibish D, Myers E, Spevak L, Compston J, Hodsman A, Shane E, Recker RR, Boskey ER, Boskey AL 2009 Use of FTIR Spectroscopic Imaging to Identify Parameters Associated With Fragility Fracture. J Bone Miner Res 24(9):1565–1571.

21. Mandair GS, Morris MD 2015 Contributions of Raman spectroscopy to the understanding of bone strength. BoneKEy Rep 4.

22. Kim K-A, Gu W, Lee I-A, Joh E-H, Kim D-H 2012 High Fat Diet-Induced Gut Microbiota Exacerbates Inflammation and Obesity in Mice via the TLR4 Signaling Pathway. PLoS One 7(10):e47713.

23. Hunt HB, Donnelly E 2016 Bone Quality Assessment Techniques: Geometric, Compositional, and Mechanical Characterization from Macroscale to Nanoscale. Clin Rev Bone Miner Metab 14(3):133–149.

24. Paschalis EP, Mendelsohn R, Boskey AL 2011 Infrared Assessment of Bone Quality: A Review. Clin Orthop Relat Res 469(8):2170–2178.

25. Zimmermann EA, Busse B, Ritchie RO 2015 The fracture mechanics of human bone: influence of disease and treatment. BoneKEy Rep 4.

26. Vijay-Kumar M, Aitken JD, Carvalho FA, Cullender TC, Mwangi S, Srinivasan S, Sitaraman SV, Knight R, Ley RE, Gewirtz AT 2010 Metabolic syndrome and altered gut microbiota in mice lacking Toll-like receptor 5. Science 328(5975):228–31.

27. Laukens D, Brinkman BM, Raes J, De Vos M, Vandenabeele P 2016 Heterogeneity of the gut microbiome in mice: guidelines for optimizing experimental design. FEMS Microbiol Rev 40(1):117–32.

28. MacGregor RR, Graziani AL 1997 Oral Administration of Antibiotics: A Rational Alternative to the Parenteral Route. Clin. Infect. Dis. 24(3):457–467.

29. Lima SF, Teixeira AGV, Higgins CH, Lima FS, Bicalho RC 2016 The upper respiratory tract microbiome and its potential role in bovine respiratory disease and otitis media. Sci. Rep. 6:29050.

30. Meyer F, Paarmann D, D’Souza M, Olson R, Glass EM, Kubal M, Paczian T, Rodriguez A, Stevens R, Wilke A, Wilkening J, Edwards RA 2008 The metagenomics RAST server - a public resource for the automatic phylogenetic and functional analysis of metagenomes. BMC Bioinformatics 9:386.

31. Wilke A, Bischof J, Gerlach W, Glass E, Harrison T, Keegan KP, Paczian T, Trimble WL, Bagchi S, Grama A, Chaterji S, Meyer F 2016 The MG-RAST metagenomics database and portal in 2015. Nucleic Acids Res 44(D1):D590–4.

32. Overbeek R, Begley T, Butler RM, Choudhuri JV, Chuang H-Y, Cohoon M, de Crécy-Lagard V, Diaz N, Disz T, Edwards R, Fonstein M, Frank ED, Gerdes S, Glass EM, Goesmann A, Hanson A, Iwata-Reuyl D, Jensen R, Jamshidi N, Krause L, Kubal M, Larsen N, Linke B, McHardy AC, Meyer F, Neuweger H, Olsen G, Olson R, Osterman A, Portnoy V, Pusch GD, Rodionov DA, Rückert C, Steiner J, Stevens R, Thiele I, Vassieva O, Ye Y, Zagnitko O, Vonstein V 2005 The Subsystems Approach to Genome Annotation and its Use in the Project to Annotate 1000 Genomes. Nucleic Acids Research 33(17):5691–5702.

33. Fierer N, Leff JW, Adams BJ, Nielsen UN, Bates ST, Lauber CL, Owens S, Gilbert JA, Wall DH, Caporaso JG 2012 Cross-biome metagenomic analyses of soil microbial communities and their functional attributes. Proc Natl Acad Sci U S A 109(52):21390–5.

34. Pereira RVV, Carroll LM, Lima S, Foditsch C, Siler JD, Bicalho RC, Warnick LD 2018 Impacts of feeding preweaned calves milk containing drug residues on the functional profile of the fecal microbiota. Sci Rep 8(1):554.

35. Goodrich JK, Di Rienzi SC, Poole AC, Koren O, Walters WA, Caporaso JG, Knight R, Ley RE 2014 Conducting a microbiome study. Cell 158(2):250–62.

36. Donnelly E, Baker SP, Boskey AL, van der Meulen MC 2006 Effects of surface roughness and maximum load on the mechanical properties of cancellous bone measured by nanoindentation. J Biomed Mater Res A 77(2):426–35.

37. Gamsjaeger S, Masic A, Roschger P, Kazanci M, Dunlop JWC, Klaushofer K, Paschalis EP, Fratzl P 2010 Cortical bone composition and orientation as a function of animal and tissue age in mice by Raman spectroscopy. Bone 47(2):392–399.

38. Kazanci M, Fratzl P, Klaushofer K, Paschalis EP 2006 Complementary Information on In Vitro Conversion of Amorphous (Precursor) Calcium Phosphate to Hydroxyapatite from Raman Microspectroscopy and Wide-Angle X-Ray Scattering. Calcif Tissue Int. 79(5):354–359.

39. Oliver WC, Pharr GM 1992 An improved technique for determining hardness and elastic modulus using load and displacement sensing indentation experiments. J Mater Res 7(6):1564– 1583.

40. Walther B, Karl JP, Booth SL, Boyaval P 2013 Menaquinones, Bacteria, and the Food Supply: The Relevance of Dairy and Fermented Food Products to Vitamin K Requirements. Adv Nutr. 4(4):463–473.

41. Nguyen TLA, Vieira-Silva S, Liston A, Raes J 2015 How informative is the mouse for human gut microbiota research? Disease Models & Mechanisms 8(1):1–16.

42. Thijssen HHW, Drittij-Reijnders MJ 1994 Vitamin K distribution in rat tissues: dietary phylloquinone is a source of tissue menaquinone-4. Br J Nutr 72(3):415–425.

43. Karl JP, Fu X, Dolnikowski GG, Saltzman E, Booth SL 2014 Quantification of phylloquinone and menaquinones in feces, serum, and food by high-performance liquid chromatography-mass spectrometry. J Chromatogr B Analyt Technol Biomed Life Sci 963:128-33.

44. Heaney RP 2001 Chapter 27 - Nutrition and Risk for Osteoporosis. In: Marcus R, Feldman D, Kelsey J (eds.) Osteoporosis (Second Edition). Academic Press, San Diego, pp 669– 700.

45. Shea MK, Booth SL 2007 Role of vitamin K in the regulation of calcification. Int Congr Ser 1297:165–178.

46. Hernandez CJ, Guss JD, Luna M, Goldring SR 2016 Links Between the Microbiome and Bone. J Bone Miner Res 31(9):1638–46.

47. Pacifici R 2018 Bone Remodeling and the Microbiome. Cold Spring Harb Perspect Med 8(4).

48. Gundberg CM, Lian JB, Booth SL 2012 Vitamin K-dependent carboxylation of osteocalcin: friend or foe? Adv Nutr 3(2):149–57.

49. Cheung AM, Tile L, Lee Y, Tomlinson G, Hawker G, Scher J, Hu H, Vieth R, Thompson L, Jamal S, Josse R 2008 Vitamin K supplementation in postmenopausal women with osteopenia (ECKO trial): a randomized controlled trial. PLoS Med 5(10):e196.

50. Morgan S, Poundarik AA, Vashishth D 2015 Do Non-collagenous Proteins Affect Skeletal Mechanical Properties? Calcif Tissue Int 97(3):281–91.

51. Poundarik AA, Diab T, Sroga GE, Ural A, Boskey AL, Gundberg CM, Vashishth D 2012 Dilatational band formation in bone. Proc Natl Acad Sci U S A 109(47):19178–83.

52. Burr DB, Akkus O 2014 Chapter 1 - Bone Morphology and Organization Basic and Applied Bone Biology. Academic Press, San Diego, pp 3–25.

53. Hunter GK, Hauschka PV, Poole AR, Rosenberg LC, Goldberg HA 1996 Nucleation and inhibition of hydroxyapatite formation by mineralized tissue proteins. Biochem J 317(Pt 1):59– 64.

54. Murshed M, Schinke T, McKee MD, Karsenty G 2004 Extracellular matrix mineralization is regulated locally; different roles of two gla-containing proteins. J Cell Bio 165(5):625–630.

55. Poundarik AA, Boskey A, Gundberg C, Vashishth D 2018 Biomolecular regulation, composition and nanoarchitecture of bone mineral. Sci Rep 8:1191.

56. Stock SR 2015 The Mineral–Collagen Interface in Bone. Calcif Tissue Int. 97(3):262– 280.

57. Yerramshetty JS, Akkus O 2008 The associations between mineral crystallinity and the mechanical properties of human cortical bone. Bone 42(3):476–82.

58. Boskey AL, Gadaleta S, Gundberg C, Doty SB, Ducy P, Karsenty G 1998 Fourier transform infrared microspectroscopic analysis of bones of osteocalcin-deficient mice provides insight into the function of osteocalcin. Bone 23(3):187–96.

59. Bailey S, Karsenty G, Gundberg C, Vashishth D 2017 Osteocalcin and osteopontin influence bone morphology and mechanical properties. Ann N Y Acad Sci 1409(1):79–84.

60. Akkus O, Adar F, Schaffler MB 2004 Age-related changes in physicochemical properties of mineral crystals are related to impaired mechanical function of cortical bone. Bone 34(3):443– 53.

61. Bi X, Patil CA, Lynch CC, Pharr GM, Mahadevan-Jansen A, Nyman JS 2011 Raman and mechanical properties correlate at whole bone- and tissue-levels in a genetic mouse model. J Biomech 44(2):297–303.

62. Yamaguchi M, Taguchi H, Hua Gao Y, Igarashi A, Tsukamoto Y 1999 Effect of vitamin K2 (menaquinone-7) in fermented soybean (natto) on bone loss in ovariectomized rats, vol. 17, pp 23–9.

63. Kim M, Na W, Sohn C 2013 Vitamin K1 (phylloquinone) and K2 (menaquinone-4) supplementation improves bone formation in a high-fat diet-induced obese mice. J Clin Biochem Nutr 53(2):108–113.

64. Okano T, Shimomura Y, Yamane M, Suhara Y, Kamao M, Sugiura M, Nakagawa K 2008 Conversion of Phylloquinone (Vitamin K1) into Menaquinone-4 (Vitamin K2) in Mice: Two possible routes fro menaquinone-4 accumulation in cerebra of mice. J Biol Chem 283(17):11270–11279.

65. Ichikawa T, Horie-Inoue K, Ikeda K, Blumberg B, Inoue S 2006 Steroid and xenobiotic receptor SXR mediates vitamin K2-activated transcription of extracellular matrix-related genes and collagen accumulation in osteoblastic cells. J Biol Chem 281(25):16927–34.

