## Supplemental Fig 1 for "The Microbial Metagenome and Tissue Composition in Mice with Microbiome-Induced Reductions in Bone Strength"

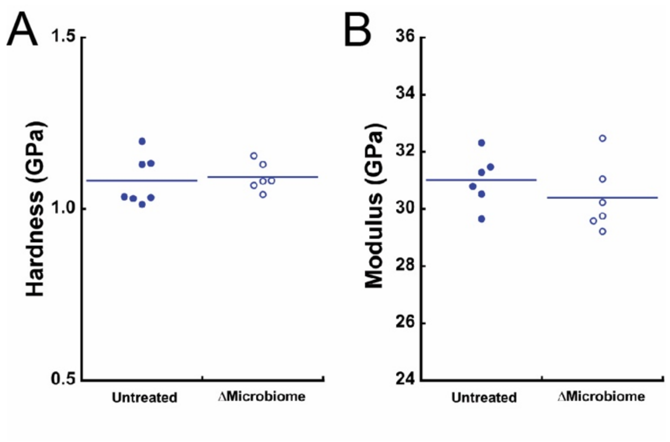


**Supplemental Figure 1.** Bone tissue material properties in the tibial diaphysis were assessed with nanoindentation. (A) Hardness and (B) modulus were similar among groups indicated no differences in compressive  material properties.
